# Collective decision making by rational individuals

**DOI:** 10.1101/363838

**Authors:** Richard P. Mann

## Abstract

The patterns and mechanisms of collective decision making in humans and animals have attracted both empirical and theoretical attention. Of particular interest has been the variety of social feedback rules, and the extent to which these behavioural rules can be explained and predicted from theories of rational estimation and decision making. However, models that aim to model the full range of social information use have incorporated *ad hoc* departures from rational decision-making theory to explain the apparent stochasticity and variability of behaviour. In this paper I develop a model of social information use and collective decision making by fully rational agents that reveals how a wide range of apparently stochastic social decision rules emerge from fundamental information asymmetries both between individuals, and between the decision-makers and the observer of those decisions. As well as showing that rational decision making is consistent with empirical observations of collective behaviour, this model makes several testable predictions about how individuals make decisions in groups, and offers a valuable perspective on how we view sources of variability in animal, and human, behaviour.

## INTRODUCTION

ollective decision making is a ubiquitous task for social animal species, including humans [1]. Whether deciding where to forage, which nest site to choose or when to move, individual decisions are greatly informed by observing the choices that others make. As recently as 2008, Ward et al. [2] were able to state that ‘little is known about the mechanisms underlying decision making in vertebrate animal groups’. Since then, however, a large literature has explored the rules governing social information use in collective decisions across various taxa, for example in insects [3], fish [2, 4], birds [5, 6] and mammals [7, 8], including primates [9] and humans [10, 11]. What links decision making in all of these groups is the presence of social reinforcement, with individuals demonstrating a strong preference for an option chosen by others, which increases with the number of others who have selected it. This reinforcement can be expressed as a social response function – the probability of selecting a given option conditioned on the number of other individuals that have previously chosen it. A large degree of variation has been observed in these social response functions, ranging from linear relationships (e.g. [3, 12]), to strongly non-linear ‘quorum’ rules [13], where the apparent attractiveness of an option appears to increase exponentially with the number of individuals choosing it, before saturating as this number passes a ‘quorum’ level. In addition to variation between taxa, studies have also highlighted how the same species can exhibit different patterns of collective behaviour under different laboratory or field conditions [14–17] highlighting the potential importance of context-dependent social responses.

Complementing these empirical studies, mathematical theories have been developed to explain why these social decision rules take the form observed. For example, Easley and Kleinberg [18] proposed a toy model for understanding collective decision making in a group of rational agents. This model, illustrated by the example of an individual selecting a restaurant to eat at, demonstrated how easily an unbreakable consensus decision could emerge, once the cumulative social information provided by past choices outweighs any new quality signal an uncommitted individual might receive. More recent work has attempted to build a fully descriptive model of such collective decision making by considering the purportedly rational beliefs and decisions of agents exposed to the social information provided by choices of others [19, 20], and the studies have been successful in reproducing the observed response functions in a variety of taxa including insects [20], fish [19–21] and birds [5, 6]. Recent extensions of these models have also considered how social responses might vary as a result of changes in environmental context [22].

However, while these models have had success in reproducing the observed features of collective decisions, this has been at the cost of internal consistency as theories of rational behaviour. An agent’s decision involves two components: (i) an estimation stage, where the focal agent forms beliefs about the quality of its options, and (ii) a decision rule, which specifies how the agent acts based on those beliefs. In the first stage the models present a broadly coherent theory of estimation based on Bayesian updating. However, beliefs are restricted to statements about whether options are ‘good’ or ‘bad’ (or ‘best’ [19]). This binary categorisation does not fully capture the range of possibilities that individuals face in expected-utility maximising, rational behaviour [23]. Choices made under uncertainty are characterised by risk-reward trade-offs, and it is not clear how these can be translated into a simple ‘good’ or ‘bad’ dichotomy, or how decisions should be made on the basis of such a classification. The second stage of these models, the decision rule, introduces further departures from rationality. Here agents are assumed to select options probabilistically based on the results of the estimation stage. Such non-deterministic behaviour is inconsistent with the idea of individuals as rational agents. This problem emerges as a result of a confusion regarding the sources of observational uncertainty in empirical studies. Because the decisions made are typically not predictable with certainty by an observer or experimenter, they are themselves deemed to be stochastic. Instead, this uncertainty can be understood by incorporating the viewpoint of the observer into the theory. The observer makes measurements of the physical and social environment that only imperfectly capture the information observed by the focal decision maker (or not observed, for example in the case of visual occlusion). From this one can recognise that the inability of the observer to predict individuals’ decisions arises from the limited access they have to the information driving those actions, not from fundamentally stochastic behaviour by the agents themselves.

Why should we be concerned about departures from rationality in these models? Given established critiques of rational-agent models [24, 25], should we not be more concerned about whether these models are consistent with empirical measurements? To this objection there are two responses. The first is that rational-agent models provide a baseline from which to measure departures from rationality. These departures are interesting because they indicate either where an evolutionary process has been unable to produce an optimal solution, or where other factors, such as cognitive cost, have produced a trade-off. Such departures can only be detected if we understand what genuinely rational behaviour looks like. The second response concerns the purported goals of previous work. One can find a reasonable empirical match to any observed social response by choosing an appropriate parametrisation of a sufficiently flexible mathematical function. It has been the explicit goal of theoretical work in this area [19, 20] to understand how these responses emerge from logical consideration of the individuals’ own estimations and actions as rational decision makers. My goal here is to fully explore the consequences of the rationality assumption to show where data can (and cannot) be explained by this fundamental principle.

This paper develops a model of collective decision making by identical, rational agents, based on three fundamental principles: (i) individuals behave as expected-utility maximising agents, based on their own beliefs about the world; (ii) each individual’s beliefs are generated from the public and private information that they have access to, using Bayesian probability updating; and (iii) empirical observation of individuals’ actions is undertaken by an observer who has their own *private* information as well as the public social information on which to base predictions and interpretations of individual behaviour. The resulting model reproduces the key successful aspects of previous research, while making additional, testable predictions about social information use that are not accounted for in existing theory.

## THEORY

Consider the classic paradigm of a group faced with a sequential, binary decision. That is, a sequence of *n* identical individuals choose between option A and option B, and can see the choices made by those ahead of them. Such a context is well approximated empirically by, for example, Y-maze experiments (e.g. [2, 26]) where individuals are asked to choose between two competing arms of a maze. I develop a mathematical framework for calculating the optimal choice for each individual, based on private information that they alone observe, and public information constituted by the observable choices made by others. I also derive mathematical expressions for the probability that an outside observer (such as an experimental scientist) will observe an individual making a particular choice, conditioned on what that observer can know about the system and the focal individual. As noted above, incorporating the observer explicitly is key to understanding the source of observational uncertainty in a fundamentally deterministic model.

### Rational choice

I start from the assumption that each of the two options each has a true utility, *U*_*A*_ for option A and *U*_*B*_ for option B. These utilities may also be understood as fitness consequences of the decision in terms of evolutionary adaptation [27]. Since the individuals are assumed to be identical, these utilities are the same for all. These true utilities are unknown, but each individual, *i*∈ 1, *…, n*, can estimate the utility of each choice based on the specific information, *I*_*i*_ that they possess. Following the rules of Bayesian-rational decision making [23], I assert that individual *i* will choose option A if and only if the expected utility of A is greater than that of B, according to *i*’s estimation. Let *x* = *U*_*A*_ – *U*_*B*_ be the true difference in utilities, then:

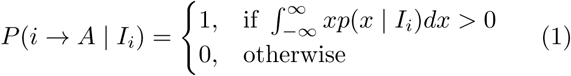

where *p*(*x* | *I*_*i*_) is a probability density representing individual *i*’s personal belief about *x*.

### Private and public information

What information does individual *i* have? I assume that *I*_*i*_ is composed of two parts, private and public. First, there is direct sensory information that *i* can perceive from the two options. For example, a foraging individual may perceive differing food odours from A and B, or a prey animal may see differing patterns of shadows that suggest one choice is more likely to lead to a predator. This is the individual’s *private* information. Secondly, if *i >* 1, then individual *i* can see the choices made by any other individual *j*, where *j < i*. This is public information – it is available to all individuals who still wait to make their choice. In common with previous work [19, 20] I make the important assumption that the choices of others provide information about the relative utilities of A and B, but do not influence the true values of these utilities. That is, an option does not become good simply because others have chosen it. In making this assumption I exclude phenomena such as predation-dilution effects [28], where the presence of conspecifics is itself desirable, or foraging competition [29], where the presence of other individuals lowers the utility of a given option.

### Prior belief

Before an individual receives any information regarding the utility difference, *x*, I assume that they have no reason to favour option A or B (any such reason should count as private information). I ascribe to them a prior belief regarding the values that *x* may take, and by symmetry centre this on zero. I further assume that by environmental habituation (either genetic or experience) they have an intrinsic idea of the scale of possible utility differences between competing choices. In this paper I will assume that prior beliefs follow a normal distribution, and without loss of generality we can measure utilities in units that set this variance of this distribution to one:

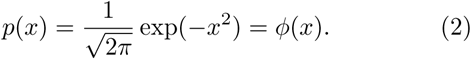

Hereafter I use *ϕ* (*x*) to refer to the standard normal distribution density function. Throughout this paper I will assume that information and expectations about the environment are normally distributed. This assumption is likely to hold well for low-level sensory information such as detecting food or predators, but may be less appropriate for more cognitively advanced tasks. The model development detailed here can be followed for any alternative distribution of interest.

### Private information

Individual *i* has access to private information that gives a noisy estimate of *x*. This may be via visual, olfactory or other sensory stimuli, but here I model this as an abstract quantity, Δ_*i*_, that is generated stochastically by the environment based on the real utility difference *x*, and a noise variance *ν*^*2*^ due to both the latent sources of environmental noise (e.g. air currents disrupting olfactory gradients) and limitations of an individual’s sensory apparatus. Mathematically, Δ_*i*_ is normally distributed with mean *x* and variance *v*^*2*^:

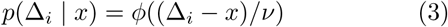

Individual *i* revises their belief about *x* in the light of their private information using Bayes’ rule:

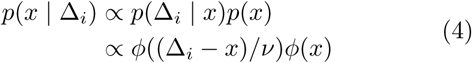

Consider the case of the first decision-maker, *i* = 1. This individual has no public, social information to draw on, and bases their estimate of *x* entirely on their private information, Δ_1_. For the purposes of making a rational decision, the important quantity for them to evaluate is the sign of the expected utility difference, 𝔼 (*x* | Δ_1_):

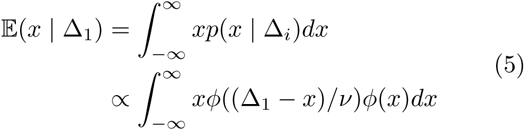

It is clear that 𝔼 (*x* | Δ_1_) > 0 *iff* Δ_1_ > 0, and therefore that the sign of individual 1’s private information dictates which option it will choose.

### Social information

Having determined that the first decision-maker uses the sign of their private information to make their choice, I now consider the case of the second and subsequent decision-makers. Individual 2 begins its estimation of *x* in the same manner as individual 1, by updating its original prior belief (which is identical for all agents), using its own private information, Δ_2_:

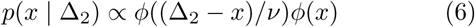

What information does the choice of individual 1 provide to individual 2? Since the choice of individual 1 does not change the true utilities of the options, its choice can only influence the estimation of individual 2 by giving information about the private information that individual 1 received. If individual 1 were to communicate its private information directly to individual 2, then the second individual could update its belief based on this new data. However, imagine that individual 1 has chosen option A. Individual 2 does not know what private information individual 1 has received, but can only infer from the observed resulting choice that Δ_1_ > 0. Therefore it must consider all possible values of the Δ_1_ that the first individual may have observed, weighted by probability, and adjust its belief accordingly. Let *C*_1_ = –1 indicate that individual 1 chose option A (and conversely *C*_1_ = − 1 for option B), then:

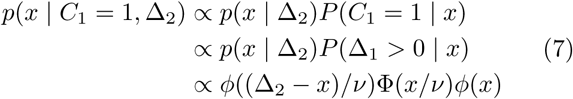

where 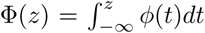 is the cumulative distribution function of the standard normal distribution. Similarly, if individual 1 had chosen B, then individual 2 would make the estimation:

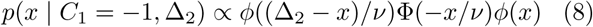

The decision of individual 2 is now governed by their expected value of *x*. Define 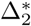 such that:

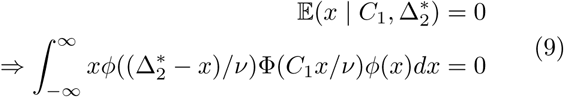

Individual 2 will now choose option A *iff* 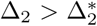, implying that 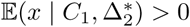.

### Subsequent decisions

To complete our view of how social information is used we need to consider the viewpoint of the third individual (assuming *n* > 2). This individual must consider not only the decisions made by individuals 1 and 2 in conjunction with its own private information, but also the order in which these decisions were made. As with all individuals, I begin by updating the universal prior, using individual 3’s private information:

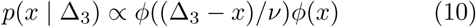

Individual 3 can also update its belief based on the choice of individual 1, in exactly the same manner as individual 2:

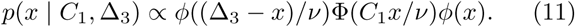

Now individual 3 needs to update its belief based on the decision made by individual 2, *C*_2_:

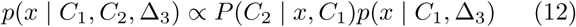

In order to evaluate the first term on the right hand side, individual 3 needs to adopt the viewpoint of individual 2 and calculate the critical value 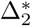 based on *C*_1_. Then, from individual 3’s perspective, the the probability of choice *C*_2_ is:

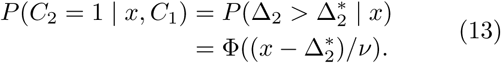

Similarly, 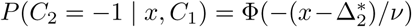. Hence, individual updates its belief based on *C*_2_ to:

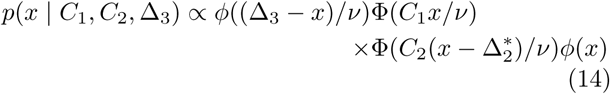

As with individual 2, we can thus evaluate a critical value,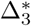, defined by:

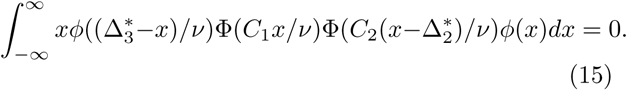

Individual 3 will now choose option A *iff* 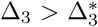. By iteratively proceeding in similar fashion we can determine the belief of individual *i*, based on its private information and the observed choices of previous individuals, *C*_1_, …, *C*_*i-*1_:

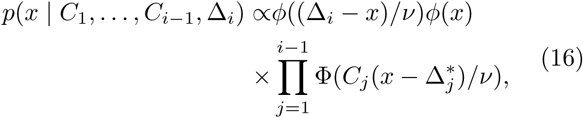

and

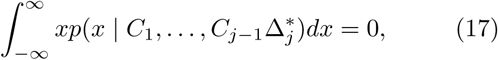

and I define 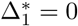.

### Observation

So far I have discussed how each individual uses private and public information to make a rational, expected-utility maximising decision. Now I consider the perspective as an observer of this process. As an observer, one is able to observe the same public, social information available to the individuals themselves – the sequence of decisions. However, the observer’s viewpoint differs in two ways. First, they have no access to the private information of any individual. Secondly, they may have knowledge about the true environmental conditions. For example, they may have designed an experiment such that *x* = *U*_*A*_ *–U*_*B*_ = 0, e.g. a Y-maze with symmetrical arms. Furthermore, especially in a laboratory setting, they may have altered the environment, such that noise levels differ from those that the individuals are habituated to.

Assume that true values of *x* and *ν*^*2*^, and also the noise variance of the experimental environment, *η* ^2^ are known. How can we calculate the probability that a individual *i* will make a specific choice? First, we can follow the calculations above for that individual, to determine the critical value 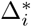, based on the observed previous decisions. We can then evaluate the probability, *conditioned on the known x and η*^2^, that individual *i*’s private information will exceed this value:

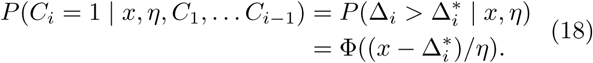

Although this equation provides the probability that the *observed* decision will be a particular option, it implies no non-deterministic behaviour; the uncertainty encoded by the probability is purely a consequence of the observer not sharing the same information as the focal decision-maker. Note that the information provided by *C*_1_, *… C*_*i-*1_ is encoded in the calculated value of 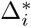. Since each critical value depends iteratively on those before, this is determined by the order of decisions made, as well as the aggregate numbers choosing A and B.

### Unordered social information

Observers not only have differing information from the individuals under observation, they also make choices about how to measure and record behaviour. As an example of this: the majority of previous studies have largely ignored the precise order of previous decisions made when measuring social responses. For comparison with this previous work, we can consider what we, the observer, would predict about the decision of individual *i*, conditioned on knowing only the number of previous individuals choosing A (*n*_*A*_) and B (*n*_*B*_). This requires us to consider the set of all possible sequences, that obey result in *n*_*A*_, *n*_*B*_, and to sum over the probability that each of these is the sequence to led to the current arrangement. This summation, combined with the decision rule for each specific sequence derived above, then gives the probability that next choice will be either A or B, conditioned on this unordered observation of social information.

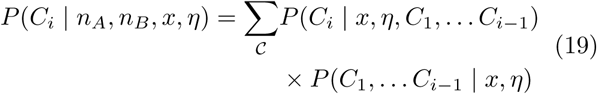

Note, this calculation assumes that the individuals themselves *are* aware of the order in which decisions were made, but that the observer has been unable to record these or has chosen not to do so.

### Conflicting information

Several experimental studies have investigated scenarios where a conflict is introduced between an individual’s private and social information, in order to identify the relative strengths of the two factors. For example, in such an experiment each individual may be trained in advance to associate food with one of two or more different colour or pattern. Individuals with different trained associations are then placed in a group and presented with a decision where each option has a colour or pattern signal (see e.g. [30]).

This scenario can be simulated by giving each agent private information drawn from a mixture of two conflicting distributions:

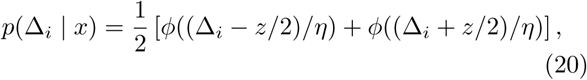

where *z* is the magnitude of the conflict in information, and **η** is again the experimental noise level. These two parameters indicate respectively how strong the training has indicated the utility difference is (e.g. the amount or quality of food provided), and the reliability of the signal (e.g. whether the food was always provided in the same quantities). With each individual’s private information drawn from this mixture distribution, a simulation of the group’s aggregate behaviour can follow as above. It should be noted here I am still assuming that the individual decision-makers have identical utility functions; the conflict between them is solely on the level of the information they have received, and not one of differing preferences.

## RESULTS

In this section I consider a variety of possible experimental and field study scenarios that illustrate the key predictions of the model.

### Role of environmental signal to noise ratio

I begin by considering an experimental field study in the habitual environment of the decision-makers. In this case the decision-maker’s previous experience gives them reliable prior information about typical signal and noise levels in their environment, while the observer can experimentally control the true utilities of possible choices. I analysed the expected behaviour when the decisionmakers are confronted with a symmetrical binary choice in which the true utility difference is zero: *U*_*A*_ *–U*_*B*_ = 0. I calculated the probability that the next decision maker would choose option A, conditioned on different previous decision sequences, and for a range of environmental noise/signal ratios. This analysis, illustrated in Figure 1 shows that environmental noise levels have little effect on the predicted choice probabilities, displaying only a slightly inflection at a ratio of one. This result can be understood intuitively by noting that higher ambient noise levels reduce the reliability of both private and public information – the decision-maker should trust its own information less, since it may result from noise,but should also recognise that the decisions of others are also more likely to be incorrect. These two effects almost perfectly balance in this analysis for a wide range of possible noise/signal ratios. Since the environmental noise/signal ratio has little effect on behaviour, I set it to a value of *v =* 1 henceforth for simplicity.

**FIG. 1.**
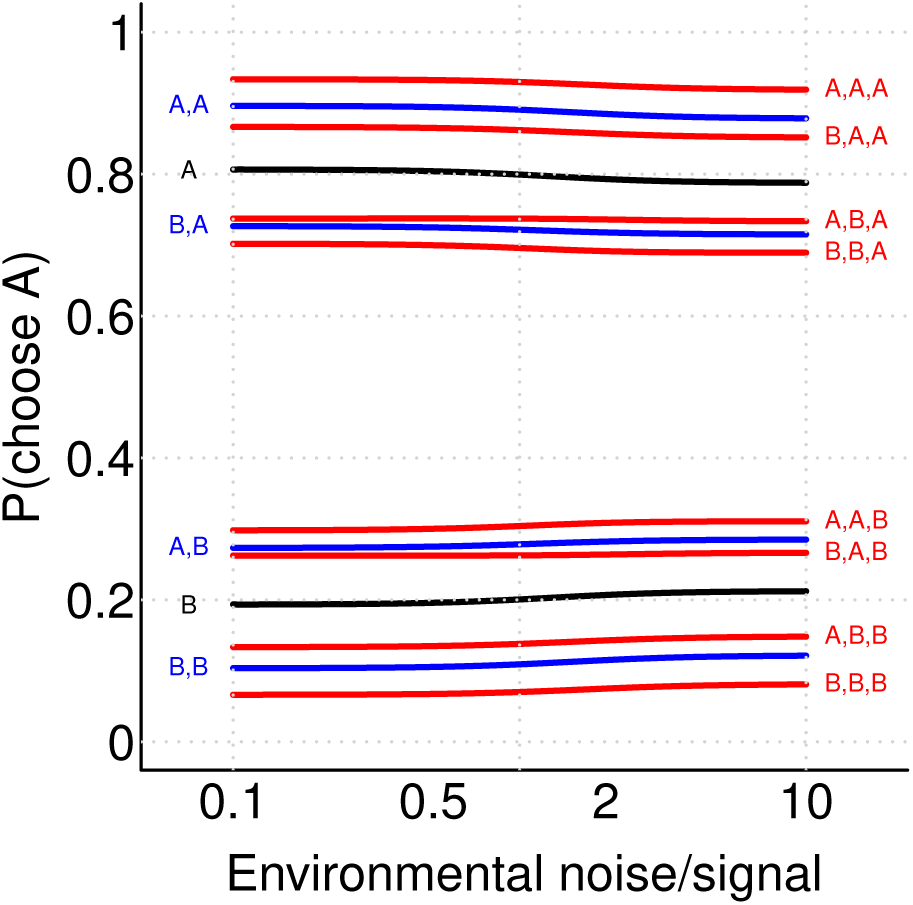
Consistency of predicted decisions across environmental noise/signal ratios, for a range of possible observed past decision sequence. Each line is labelled with the corresponding sequence of past decision. The black line shows cases with one previous decision maker, blue lines two and red lines three respectively.

### Observed social interaction rules

My model gives the probability that a focal individual will make a given choice conditioned on any ordered sequence of previous decisions. However, in practice researchers are often either unable to observe this precise sequence, or choose to ignore the details of the order in which decisions were made, focusing instead only on the number of individuals who have previously chosen A or B in aggregate. To make comparisons between the theory developed here and previous work, I therefore calculated the expected observations on this aggregated level by considering all possible sequences of previous choices that could have led to an aggregate state *n*_*A*_, *n*_*B*_ (see equation 19). To illustrate the predicted observations a researcher would make in such an experiment, we consider two hypothetical experiments, each with ten individuals and in which the ambient noise level matches that of the habitual environment. The first experiment uses symmetric options (*x* = *U*_*A*_ *–U*_*B*_ = 0) The predicted observations made in these experiments are shown in Figure 2. Panel A shows the probability that a focal decision maker will choose option A, conditioned on known *n*_*A*_ and *n*_*B*_. Contour lines of equal probability show a radial pattern that is suggestive of Weber’s Law of relative differences [31]. To illustrate this further, panel B shows how the probability of choosing option A varies with the relative proportion of previous decisions: (*n*_*A*_)*/*(*n*_*A*_ + *n*_*B*_). We see that the results averaged over all possible sequences (red points) show a very close linear trend. However, this apparently simple relationship results from the weighted average of sequence-specific probabilities, shown with black points, where larger points indicate more probable sequences. Evaluating the probability of generating each possible sequence of decisions, we can determine the probability for the final value of *n*_*A*_ after all decisions have been made (panel C). This exhibits the classic Ushaped distribution that is characteristic of observations in many collective decision-making studies (e.g [13]). At first glance this result conflicts with the pattern in panel B – a sequence of decisions made according to Weber’s Law would be expected to results in a uniform distribution equal probabilites for final values of *n*_*A*_ (see [32]). This apparent contradiction is resolved by noting that the linear relationship in panel B does not hold as a decision rule in its own right, but only as the average behaviour aggregated over many possible sequences of decisions using the true behavioural rules shown in equations 16 and This highlights how apparently straightforward analysis of empirical data may lead to erroneous conclusions about underlying behavioural mechanisms.

**FIG. 2.**
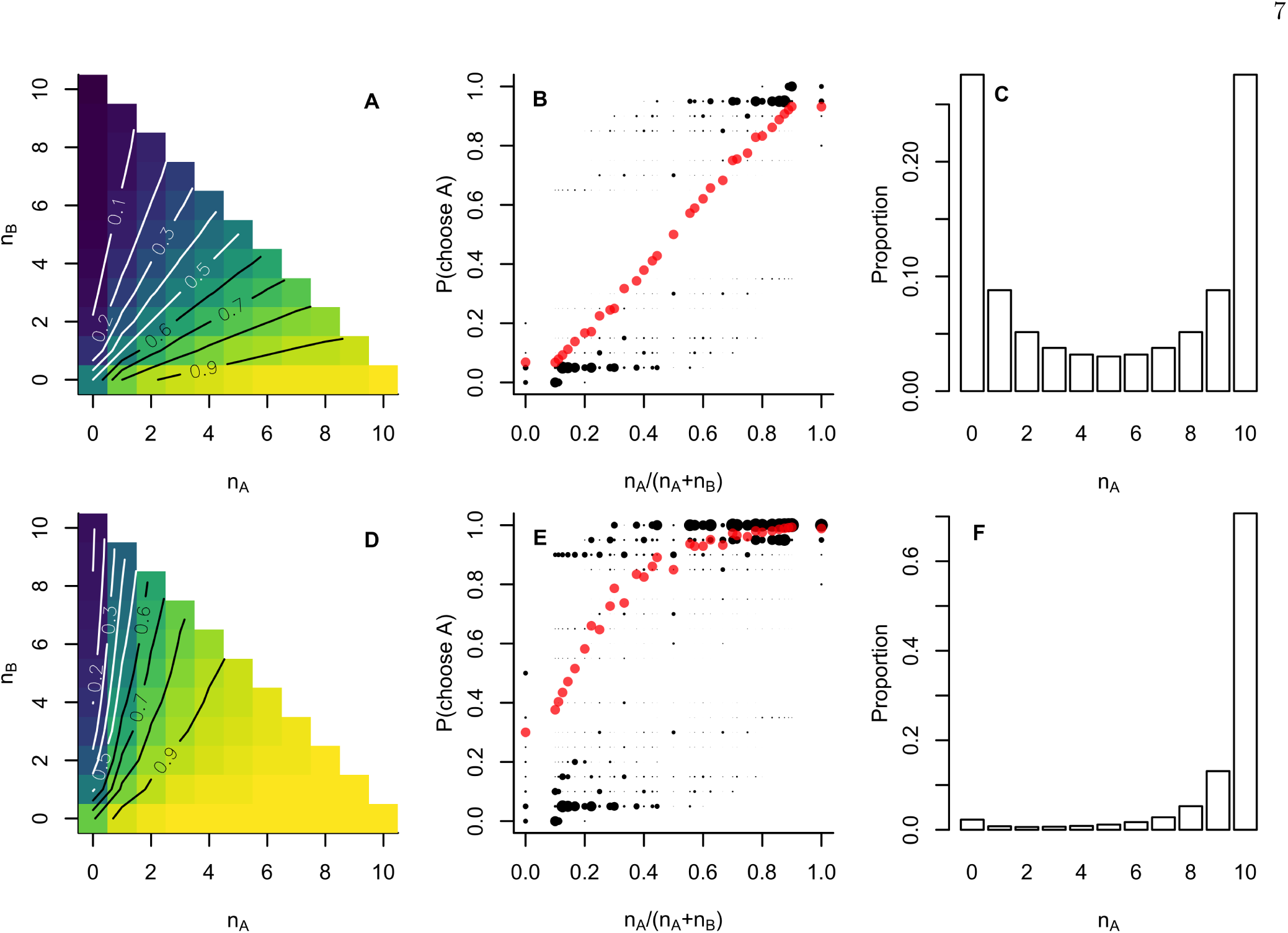
Results of a two hypothetical studies of collective decision making. Panels A-C show predicted observations made from a symmetric experimental setup (*x* = 0), while panels D-F show those predicted from an experiment in which option A is superior by an amount typical in the habitual environment 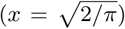: (A, D) the probability, from the perspective of an external observer, that a focal agent will choose option A, conditioned on the number of agents, *nA, nB*, previously choosing options A and B respectively; (B, E) the probability of the focal agent selecting option A against the proportion of previous agents selecting A; red points indicate the average across all possible sequences of previous choices, while black points indicate probabilities conditioned on specific sequences (discretised to intervals of 0.05), with larger points indicating more likely sequences. The average trend shows the relationship that would be observed in an experiment where sequence information was discarded; (C, F) the probability of possible aggregate outcomes, defined as the number of agents in total that will select option A, showing the high probability of consensus decisions, and of collectively choosing the higher-utility option in the asymmetric scenario.

In addition to a symmetric experimental setup, I also consider a hypothetical experiment in which one option is objectively better than the other, for example through the presence of food (e.g. [33]), or the absence of a predator (e.g. [26]). In this example I assume that option A is better, and set 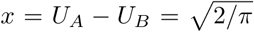. This value is equivalent to the average absolute difference in utilities in the habitual environment, and thus represents a ‘typical’ decision for the agents to make. I made predictions of the observed decisions made by ten agents as in the symmetric case, the results of which are shown in Figure 2 D-F. In this case we see that, as expected, decisions systematically favour the higher utility option. As above, the decisions observed as a function of *n*_*A*_ and *n*_*B*_ hide a broader variety of social contexts defined by the ordering of decisions (panel E).

### Context specificity

The hypothetical experiment above was assumed to take place in an environment where noise levels were the same as the decision-makers’ habitual experience. I showed that this habitual noise level did not in itself have a strong influence on predicted decisions – individuals habituated to noisy environments should be no more or less likely follow one another *in their own environment* than those from less noisy habitats. But what if individuals are removed from their own habitat and placed in an unfamiliar environment? As an example, consider collecting fish which usually shoal in somewhat murky and strongly odoured rivers or lakes, and placing these in clear water in a uniform, plastic experimental arena (e.g [13, 26, 34–36] and many others). What impact might this change have on their behaviour? One possibility is that a severe change of environment may lead to erratic or pathological behaviour as a result of distress or disorientation, which I do not account for here. Another possibility is that the fish continue to follow social interaction rules that have evolved to be near-optimal in their own habitat, without accounting for the changed context.

In the relatively sterile laboratory conditions, the ambient levels of noise such as stray odours may be far lower than in the wild. In combination with a symmetrical experimental setup as above (*U*_*A*_ = *U*_*B*_), this means that an individual is less likely to observe private information of a sufficient magnitude to contradict the apparent social information provided by its conspecifics, and will therefore be more inclined to aggregate with and follow these other individuals than it would in the wild. Conversely, a strongly-lit laboratory environment may introduce a greater intensity of visual noise simply by virtue of the greater overall intensity of visual stimulation. I explicitly calculate the effect that experimental noise has using equation 18, varying the ratio between experimental and habitual noise levels. That is, I assume that the agents continue to act rationally in the belief that noise levels remain at those their habitual environment, while in fact the noise levels depart from this baseline. As shown in Figure 3, I find as expected that lower experimental noise levels increase the tendency to follow the majority (panels A, B), resulting in a greater aggregate consensus (panels E, F). Higher noise levels reduce the weight of social information (panels C, D) and thus prevent consensus from emerging (panel G, H). Conversely this means that for a given experimental noise level, individuals from noisier habitual environments are expected to aggregate more strongly and behave more socially than those from habitats with less noise.

**FIG. 3.**
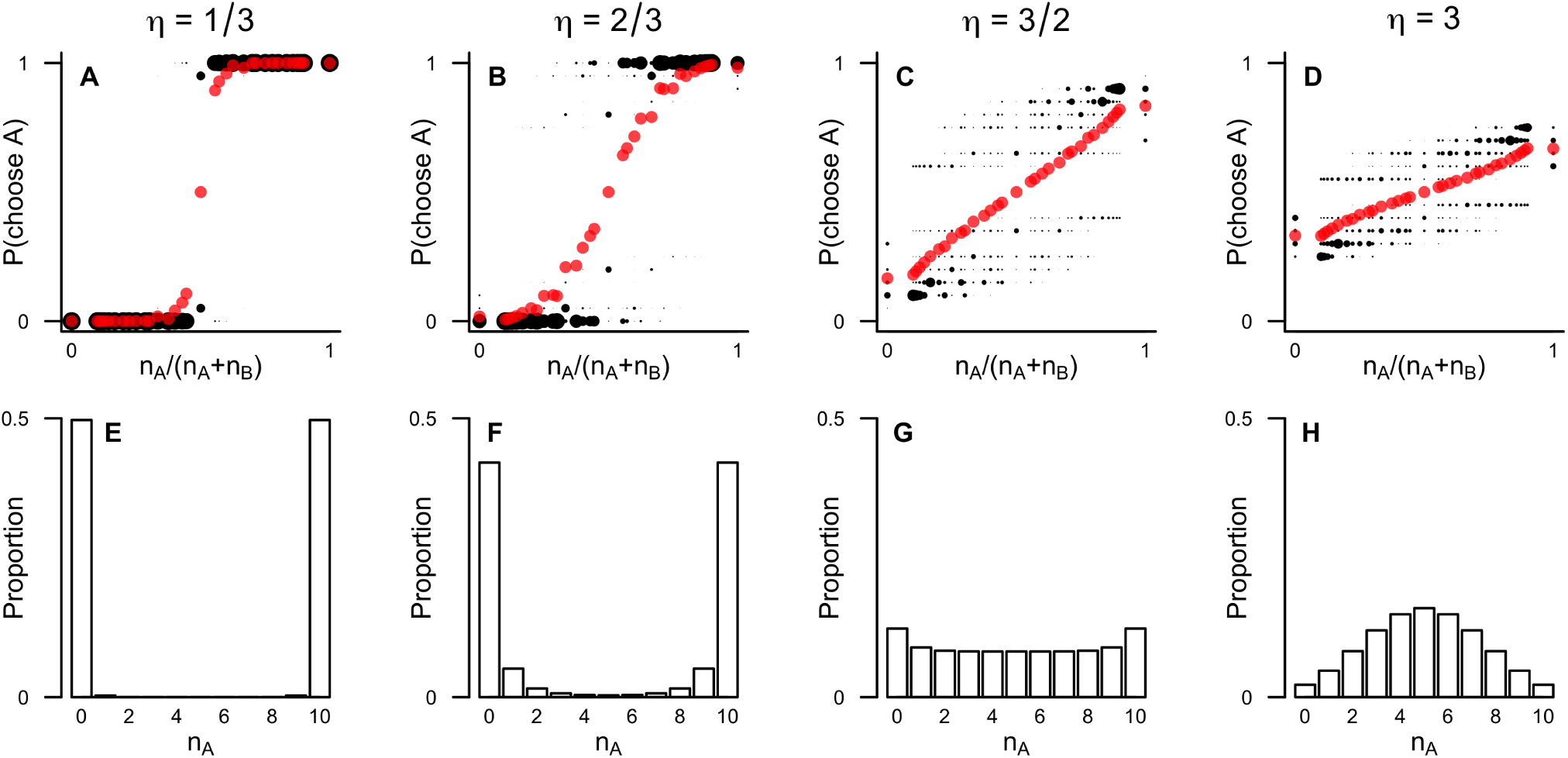
The effect of varying the experimental noise level. Panels A-D show the social response function for hypothetical experiments with noise levels in the ratios **η** = 1*/*3, 2*/*3, 3*/*2 and 3 relative to the habitual environment. As in Figure 2, red points indicate the average observed response, while black points indicate specific sequences of past decisions, with size indicating the relative probability of each sequence. Panels E-H show the corresponding aggregate outcome for each experimental noise level. Social response and aggregate cohesion is stronger in experiments with noise levels lower than the habitual environment (**η* <* 1), and correspondingly weaker in experiments with greater noise levels (**η* >* 1).

### Dynamic social information

As noted previously, the predictions my model makes about the decision a focal individual will make, conditioned on the available public information, depends strongly on the precise order in which previous decisions were made. To investigate this further, I now focus on the relative importance of the most recent decision in particular. Using equations 16, 17 and 18, I calculated the probability, in a symmetric experimental setup, that a focal individual will choose option A, conditioned on previous sequences of decisions of the form *C* = {–1,*–*1, …, 1}. That is, sequences in which the most recent previous decision was option A, after a series of individuals choosing B. In most models such sequences with many individuals choosing B and only one choosing A – would result in a high probability that the focal individual would choose B, following the majority. By contrast, as shown in figure 4, I predict that the focal individual is most likely to choose A, regardless on the weight of the majority for B. While longer series of earlier choices for B do make the probability to select A somewhat lower, this nonetheless always remains above 0.5. This result might initially appear counter-intuitive: why should the social information provided by a large majority of other individuals be outweighed by one recent decision? However, this neatly illustrates the consequences of taking seriously the idea of identical, rational agents. The focal individual, observing the most recent decision, must conclude that the individual making that decision has observed private information which is sufficient to outweigh all the previous public social information. The focal individual cannot observe this private information directly, but since the agents are identical it can infer that had it seen this information itself, it would have made the same decision. Therefore it must conclude, prior to observing its own private information, that the available public information is now in favour of A. Since, in a symmetric implementation of the model with *U*_*A*_ = *U*_*B*_, the focal individual’s private information is equally likely to favour either choice, the observational prediction is that the most-recent decision will be followed on the majority of occasions, regardless of the overall number of previous choices made for either option.

**FIG. 4.**
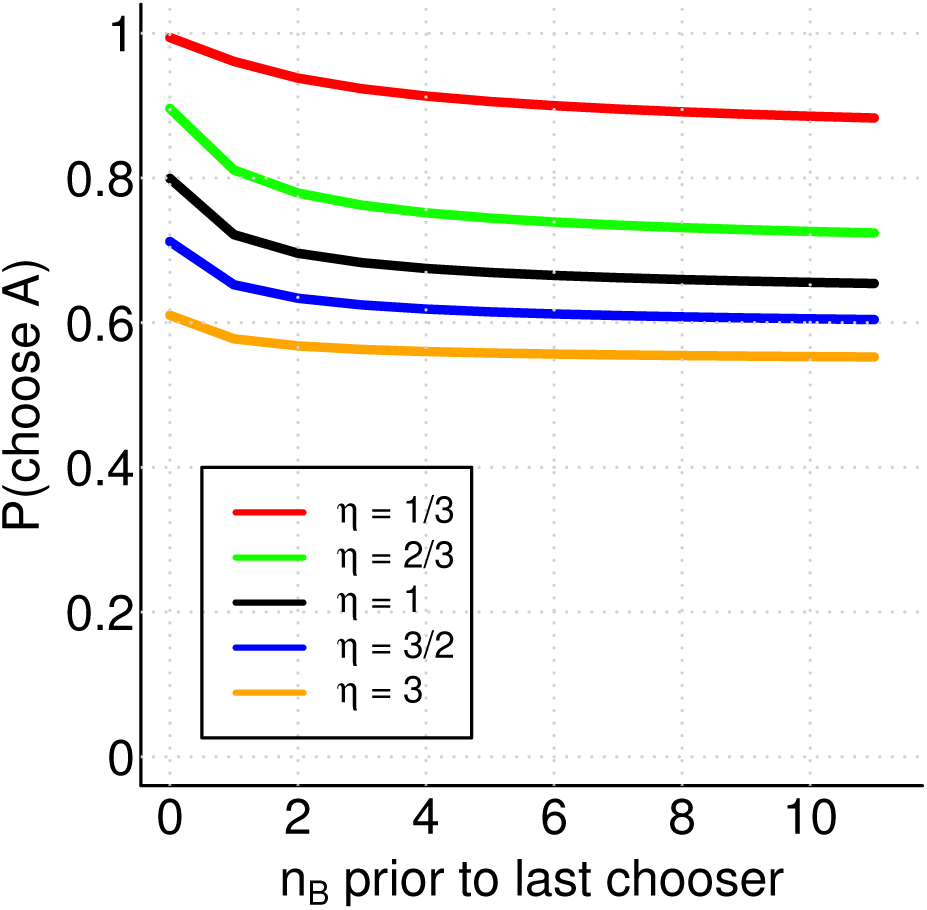
Predicted probability for a focal individual to choose option A, conditioned a sequence of past decisions of the form: B,…, B,A, evaluated for a range of experimental to habitual noise ratios (labelled by line). In all cases the probability to choose A remains above 0.5, regardless of the size of the majority choosing B.

### Conflicting information

I considered a scenario of 10 individuals that receive conflicting private information according to equation 20. I evaluated the probability of each possible aggregate outcome at each of twenty different magnitudes of conflict between *z* = 0 and *z* = 10, and five different experimental noise levels of **η** = 1*/*3, 2*/*3, 1, 3*/*2 and 3. The results, shown in Figure 5, show the degree of group consensus for each scenario, between zero (individuals split between two options equally) and one (all individuals choosing the same option). As already shown above, consensus is not guaranteed even when conflict is zero, and noisier experimental setups tend to reduce consensus. Increasing the magnitude of conflict decreases the expected degree of consensus. With sufficient conflict consensus breaks down entirely and each individual simply follows its own private information, leading to consensus values in line with those expected from a binomial distribution (dashed line). For high noise conditions the decline of consensus is gradual from an initially low value, whereas in low noise conditions there is a clearer transition from consensus to independent decision making. When training information is highly reliable compared to that found in the habitual environment (*η* = 1*/*3) this transition is very sharp, and the intermediate range between full consensus and completely independent decisions is very narrow.

**FIG. 5.**
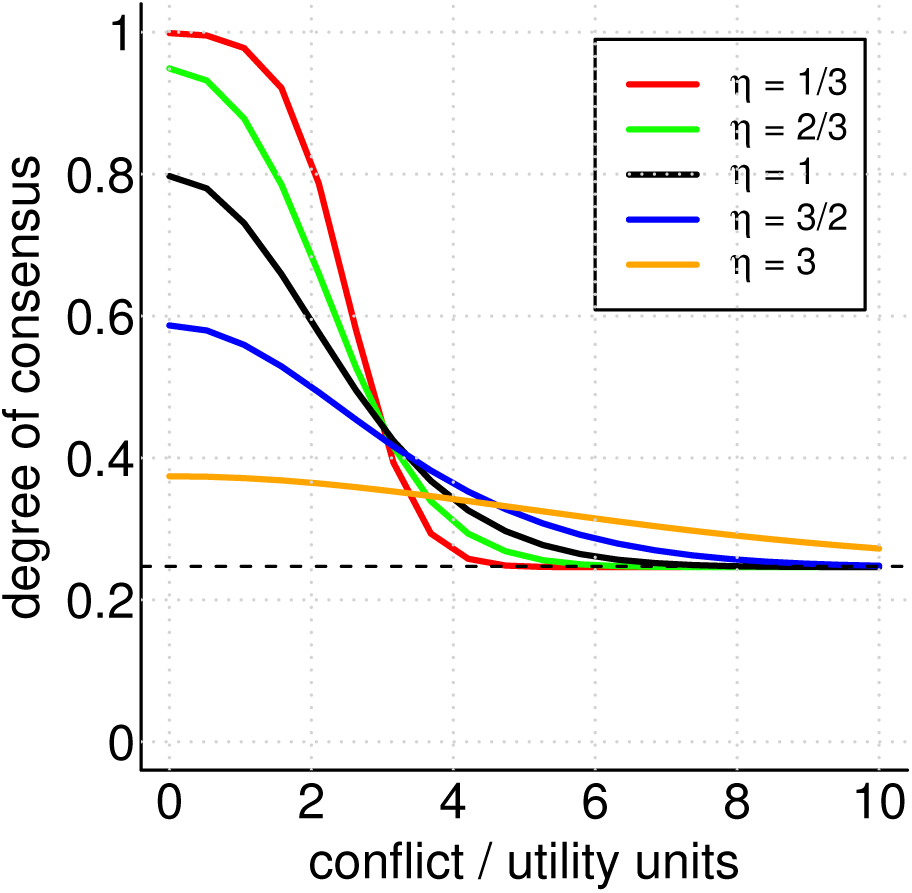
The degree of consensus in groups when individuals receive conflicting information, as a function of the magnitude of information conflict and the experimental noise level (*η*). The dashed line shows the expectation from a binomial distribution where each individual chooses independently

## DISCUSSION

I have developed a model of collective animal decision making based on perfectly rational individual decisions by identical individuals in the context of private and public information. Using this model I have explored the consequences of rational decision making, from the perspective of an observer who also has only partial information about individuals under observation. The results shown here demonstrate fundamental similarities both to earlier models of collective decision making, and to the key features of observed behaviour across a variety of taxa [20].

Formulating this model focused attention on the under-appreciated role of the observer and experimental context in understanding why animals under study make particular observed choices. Specifically, I have shown that even when agents themselves are purely rational (and therefore act deterministically on their own information), their actions appear random to the observer as a result of the agents’ private information. Furthermore, the observer potentially influences both the behaviour of the individuals under study (through their control of experimental conditions) and the interpretation of behaviour observed (through their choice of what to measure). Both of these aspects of observation are under-appreciated in the collective behaviour literature, and are potentially responsible for a substantial proportion of the variance in empirical observations. It should be noted that this perspective does not imply that the actual process being observed is dependent on the observer. Two different observers making measurements of the same experiment will observe the same decisions being made, but they may come to different conclusions depending on what they know about the experimental setup, and what they choose to measure.

This model predicts that the decision-making process is context specific. To the degree that laboratory conditions represent a lower noise environment than the wild, I anticipate that observed social tendencies will be more pronounced than in the wild. In human behaviour, this offers an explanation for why individuals exposed to social information in laboratory experiments exhibit stronger effect sizes than those exposed in more naturalistic environments [16]; the laboratory environment is subject to less spurious information that can contradict the social information presented. A recent study of contextdependent collective behaviour in sticklebacks also found that these fish were more cohesive in featureless environments than those with more distractions such as food or plant cover. The model also offers an ecological explanation for the differing social behaviours observed in different species in similar contexts. For example, Aron *et al.* [37] found that Argentine ants (*L. humile*) showed a stronger preference for social information over private information compared to garden ants (*L. niger*), and reflected that this may be explained by the differing ecology of these two species: Argentine ants are restless migrators feeding on novel food sources (a high noise/signal environment), while garden ants are sedentary and feed on well-established food sources (a low noise/signal environment). In similar laboratory conditions, *L. humile* therefore arguably experiences a greater reduction in noise relative to its habitual environment, leading to a prediction of stronger social behaviour. Similarly, Wright *et al.* [38] found that wild-strain zebrafish exhibited a stronger shoaling tendency than laboratory-strain specimens when both were tested in the same laboratory environment. This was attributed to differences in predation risk but could also reflect informational differences in each strain’s habitual environment.

One should, therefore, be careful when interpreting differences in laboratory behaviour between species from different environments, as this may betoken differing contrasts between the laboratory and the wild, rather than different habitual levels of sociality. With the development of increasingly advanced tracking technology, researchers have recently re-oriented towards studying animal behaviour in the wild [39–41], and to studying human social behaviour outside of the laboratory [42–44]. This study supports this trend; collective behaviour in the wild may vary significantly from that in the lab, and understanding natural behaviour thus requires studying the animals in their habitual environment. An interesting corollary of this finding is that social behaviour may be expected to change and become more apparently rational in the laboratory over time in cognitively-plastic species such as humans, as they habituate to the new environment. Indicative results of such an effect in a related domain have been shown for example by Burton-Chellew, Nax & West [45], who found that initially ‘irrational’ pro-social behaviour by players in a public goods game become more ‘rationally’ self-serving as the game was repeated many times under the same laboratory conditions. Whether or not similar plasticity is seen in the laboratory use of social information is worthy of further study.

I also investigated how context specificity affects behaviour when individuals are given conflicting information, for example through training prior to the experiment. I found that in laboratory conditions where complete consensus decision making is the norm (low noise relative to the habitual environment), the reaction of individuals to conflicting information is predicted to be strongly non-linear with respect to the magnitude of the conflict, with a sharp transition between consensus and independent decision making. However, this transition, which might be observed as a critical threshold in the laboratory, would be less clearly observed in more natural conditions (*η* = 1), where the model predicts a more gradual decline in the degree of consensus achieved as the magnitude of conflict is increased. Again, this highlights how behaviours observed in laboratory experiments may not be directly translatable to wild behaviour. Furthermore, I have shown that collective decision making with conflicting information depends on both the magnitude and reliability of the information individuals receive, whereas previous studies have often treated these distinct informational features ambiguously in force-based models of collective decision making (e.g. [46, 47]).

Comparing the model predictions and experimental studies highlighted a further important data analysis consideration. I measured how individual decisions varied in the context of how many others had previously chosen different options, but without any information about the sequence of those choices. In the symmetric case, agents exhibited an apparent social interaction that depended linearly on the number of other individuals choosing either option – a Weber’s law response function. The linearity of this relationship is apparently at odds with the strong tendency to consensus at the aggregate level. Perna *et al.* noted the same apparent conflict in their experimental study of trail formation in ants [3]. They proposed that the conflict could be resolved through the introduction of stochastic noise in the decision-making process. In contrast, I have shown that this conflict can be be resolved within a rational model by focusing on the importance of ordering in the sequence of previous decisions. The sequence of decisions has often been ignored in previous work or relegated to additional material not central to the study’s key insights; this model forces us to recognise the central place that ordering has in understanding rational social behaviour. As shown by Perna, Gregoire & Mann [48], inferred social responses depend intimately on how social context and behaviour are measured. The results shown here should reiterate that so-called ‘model-free’ data-driven analysis [49] is an illusion – even when no model is specified, it is implicit in the choices made by researchers regarding what to measure.

Looking more closely at the effect of ordering also revealed testable predictions about the use of social information for which few existing data are available. Specifically, the model predicts that social information associated with the most recent decision makers should have an overwhelming impact on the focal agent. Excluding their own private information, a focal agent should always conclude that the total social information favours the most recent decision made, precisely because they must believe that, as identical rational agents, they would have made the same decision given the same information. From an observational perspective this means that we should expect to see the most recent decision being more predictive of the next decision than the aggregate numbers of previous choices. To the best of my knowledge no experiment has previously tested this specific hypothesis. However, some suggestive evidence of such an effect has been seen in at least one previous study [50], where decisions by humbug damselfish to switch between two coral regions were best predicted by the most recent movement of a conspecific. Preferential following of recent decision makers may also help to explain the sensitivity of groups to changes of movement by relatively few initiators and the corresponding prevalence of ‘false alarms’ in groups of prey animals [51]. However, reliably inferring such a behavioural rule from observational data is difficult – potentially unknown information driving the most recent decision-makers choice may also be influencing the focal individual. The prediction that the most recent decisions provide the most salient social information could be tested experimentally by inducing a conflict between the most-recently observable decision and the majority of previous decisions, for example through the use of artificial substitute conspecifics (e.g. [52, 53]).

Real-world animal species, including humans, only approximate perfect rationality [24, 54] and typically only in contexts in which adaptation has taken place. A model of rational behaviour should not be mistaken for a detailed understanding of biological cognition: behaviour results from biological processes that are subject to evolutionary pressure and physical constraints, and understanding the these biological mechanisms will be important in gaining further understanding of how animals cognitively represent and process social information [55]. Nonetheless, the results of this study serve an important purpose. Assumptions of rationality and optimality are an important tool in understanding adaptive behaviour. These assumptions, when posited [19, 20], should be followed to their logical conclusion. Otherwise, it is impossible to determine, by comparing predictions and empirical data, whether or not observed behaviour supports them. Precisely because departures from rationality are of such profound interest to biologists, economists and psychologists, it is important to be precise in identifying what those departures are.

